# Alcohol consumption drives sex- and region- specific disruption of somatostatin signaling in mice

**DOI:** 10.1101/2025.06.16.659907

**Authors:** Dakota F. Brockway, Keith R. Griffith, Andrew J. Kacala, Lauren Bellfy, Laurel R. Seemiller, Matthew Ulrich, Joseph M. Ricotta, Md Shakhawat Hossain, Janine L. Kwapis, Patrick J. Drew, Nicole A. Crowley

## Abstract

The prefrontal cortex (PFC), which is thought to be disrupted early in the cycle of substance use and addiction [1], is comprised of a complex microcircuit of long-range glutamatergic pyramidal neurons controlled by GABAergic-expressing local inhibitory neurons [2, 3]. Somatostatin (SST)-expressing neurons are a subpopulation of these local GABAergic inhibitory cells and provide both peptidergic and GABAergic control over these PFC circuits [3, 4], and are disturbed following alcohol consumption in humans [5] and in rodent models [6, 7]. However, little is known about how endogenous SST peptide signaling is affected by alcohol. Using *ex vivo* electrophysiology, immunohistochemistry, *in situ* hybridization, and behavior, we demonstrate robust down-regulation of SST control over pyramidal output activity in the prelimbic (PL), but not infralimbic (IL), PFC after alcohol exposure. We also show this is likely mediated by changes in SST receptor expression levels and not disrupted expression or capacity for release of SST peptide, suggesting postsynaptic homeostatic changes to SST signaling following binge alcohol consumption in mice that may underlie post-alcohol dysregulation in mood. This provides insight into how voluntary alcohol consumption disrupts PFC peptide signaling and suggests a potential therapeutic target for the treatment of alcohol use disorder (AUD).

## INTRODUCTION

Alcohol consumption affects a variety of brain regions including the striatum, amygdala, thalamus, and hippocampus, with particularly marked disruption to the prefrontal cortex [1, 8–10]. The prefrontal cortex (PFC) provides top-down control over complex behaviors, such as those requiring integration of sensory stimuli and perception, internal states, memory, motivation, and emotions [11, 12], and is known to regulate both substance use broadly, and binge drinking behavior specifically [1, 6, 13]. Converging evidence indicates that pyramidal neurons in the prelimbic (PL) subregion of the PFC are greatly disrupted in multiple rodent models of alcohol exposure, including vaporized ethanol exposure [14, 15], forced abstinence following two bottle choice [16], and drinking-in-the-dark models of binge ethanol consumption [6]. Work across multiple labs and drinking models consistently points to a hyperexcitable phenotype of PL cortex pyramidal neurons following alcohol exposure [6, 15], and chemogenetic inhibition of pyramidal neurons in the PL cortex reduces drinking. Pyramidal cells in the PL cortex are therefore both causally involved in controlling drinking behavior and are also sensitive to the effects of binge drinking. Despite the strong evidence that PL cortex pyramidal neurons themselves are affected by binge drinking, we lack a mechanistic cellular understanding of why these changes occur, and importantly, how changes in other cells can drive pyramidal neuron changes.

Recent work by both our lab [6, 17] and others [2, 7, 18, 19] highlights the importance of somatostatin (SST) neurons within the PL cortex as a major regulator of PFC output [2, 3, 20]. New findings in humans also corroborates the general role of SST neurons in substance misuse, finding that SST gene expression modulates alcohol-induced changes in functional connectivity in healthy adult men [21], and that SST peptide levels are associated with alcohol dependence, with lower SST expression corresponding with greater dependence [5]. The SST peptide itself, which is a crucial modulator of PFC circuits [3, 4, 22, 23] and a promising therapeutic target [24] may be directly affected by binge drinking, but little is known about how, and further investigation of this peptide and its interaction with alcohol consumption is imperative.

Here we investigated how a mouse model of binge drinking leads to adaptations of SST signaling in the PFC. Using slice electrophysiology, immunohistochemistry, *in situ* hybridization, and optogenetics, we demonstrate alcohol-induced deficits in SST function in the prelimbic (PL), but not infralimbic (IL), subregions of the PFC. This work provides a putative mechanism by which binge drinking disrupts SST engagement of PFC cortical circuits and leads to PFC hyperexcitability via deficits in receptor-mediated inhibition, and presents the conceptual groundwork for a novel therapeutic approaches for the treatment of AUD.

## MATERIALS AND METHODS

### Animals

All animal procedures were performed in accordance with the Institutional Animal Care and Use Committee (IACUC) at The Pennsylvania State University. Adult (over 8 weeks of age) male and female C57BL/6J mice (stock #000664, The Jackson Laboratory), hemizygous SST-IRES-Cre mice (stock #013044, The Jackson Laboratory) Ai32 mice (stock # 024109, The Jackson Laboratory), and Ai9 reporter mice (stock #007909, The Jackson Laboratory) on a C57BL/6J background were bred in-house. All mice were single-housed (vivarium temperature 21°C, ±1°C) on a 12 hr reverse light cycle (lights off at 7:00 am) at least one week before experimental manipulation and for the duration of the experiments, as is consistent with choice alcohol consumption paradigms [25]. Mice had *ad libitum* access to food and water (except for during the DID procedure, described below, when water was removed for a short period of time).

### Drinking in the Dark

Drinking in the Dark (DID) was conducted as previously published [25, 26]. Mice received 20% (v/v) ethanol (EtOH; Koptec, Decon Labs, King of Prussia, PA) in tap water, 3 hr into the dark cycle for 2 hr (i.e., 10 a.m. to 12 p.m.) on three sequential days. On the fourth day, they received EtOH for 4 hr (i.e., 10 a.m. to 2 p.m.). Following the binge day, mice had three days of abstinence before repeating the cycle 3 more times (4 cycles total). For experiments involving surgical manipulations, mice underwent one week of DID before surgeries [27].

### Behavior

The Open Field Test (OFT) and Elevated Plus Maze test (EPM) were conducted in a subset of male and female mice at 24 hr post-DID to confirm the emergence of deficits in exploratory and anxiety-like behaviors. Detailed descriptions of the behavior are available in the ***supplementary materials***. Behavior was analyzed with Deep Lab Cut as previously published [4, 28].

### Drugs

SST (Bachem, H-1490) was dissolved in ddH2O at 1 mM, aliquoted at 50 µL, stored at –20°C, and diluted to 1 µM in aCSF (artificial cerebrospinal fluid; described in more detail below). Tetrodotoxin (TTX) (Abcam, ab120054) was dissolved in ddH2O at 5 mM, aliquoted at 50 µL, stored at –20°C, and diluted to 500 nM in aCSF. 3 mM Kynurenic acid (Sigma, K3376), 25 μM picrotoxin (Hello Bio, HB0506), and 1μM CGP 55845 (Tocris, 1248) was added to the aCSF for select experiments.

### Enzyme immunoassay for photostimulated release of SST

Acute brain slices for photostimulation were prepared as previously described and as detailed in the ***supplementary materials*** [4, 22, 29].

### Patch clamp electrophysiology

Mice were deeply anesthetized via inhaled isoflurane (5% in oxygen, v/v) and rapidly decapitated. Brains were quickly removed and processed according to the N-methyl-D-glucamine (NMDG) protective recovery method [30] (see ***supplementary materials*** for detailed methods). Drugs were included in the aCSF as described per experiment. Pyramidal neurons in layer 2/3 of the PL or IL cortex were identified by location from midline, morphology (prominent triangular soma and apical dendrites for pyramidal neurons), and membrane characteristics, consistent with previously published electrophysiology in PL cortex layer 2/3 pyramidal neurons [6, 18]. All experiments used a potassium-gluconate (KGluc)-based intracellular recording solution, containing the following (in mM): 135 K-Gluc, 5 NaCl, 2 MgCl2, 10 HEPES, 0.6 EGTA, 4 Na2ATP, and 0.4 Na2GTP (287–290 mOsm, pH 7.35). For all experiments where drugs were added to the aCSF, slices were perfused for a minimum of 10 min with aCSF containing the drugs, and slices were discarded after each experiment. Input resistance was monitored continuously throughout the experiment, and when it deviated by more than 20% the experiment was discarded.

### Somatostatin immunohistochemistry and in situ hybridization (ISH; RNAscope)

Adult (over 8 weeks of age) C57BL/6J mice were used for immunohistochemistry experiments. 24 hr following 4 cycles of the DID protocol, mice were processed for immunohistochemistry (see ***supplementary materials*** for detailed methods). 5-8 sections of the dorsal PFC were quantified and averaged to obtain one value per mouse. Imaging, quantification, and analysis was done blinded to condition and sex.

RNAscope was performed on fresh-frozen tissue as previously published [31] using the RNAscope Multiplex Fluorescent v2 Assay kit (see ***supplementary methods*** for extended protocol). The RNAscope Probe-Mm-SSTR4-C2 (ACDBio, #416641-C2) and DAPI were used to stain the PL subregion of PFC. The probe was visualized with Opal 690 (Akoya Biosciences, #FP1497001KT). Slides were imaged with a Leica STELLARIS 5 confocal (Leica). Colocalization of DAPI regions and SST receptor (SSTR) puncta was calculated using a custom semi-automated algorithm in ImageJ. In brief, RNAscope images were thresholded and binarized to locate individual DAPI regions and SSTR puncta. DAPI regions with any detectable overlap of SSTR puncta were graded as SSTR+, and the fraction of SSTR overlap was used to estimate transcript density for each DAPI region. Cell density and SSTR+ cell prevalence were quantified as the number of identified DAPI regions per image and the fraction of cells expressing SSTR regions per image, respectively. Transcript density in SSTR+ cells were estimated as the fraction of DAPI overlap with any SSTR puncta.

### Statistics and data analysis

All statistics, data analysis, and figure preparation were performed using MATLAB (MathWorks), GraphPad Prism 7.0, and Adobe Illustrator. All data are represented as mean ± SEM. Statistical significance was set as *p* < .05. A 2-way ANOVA (factors: sex, DID condition) was used for SST cell counts, SST immunofluorescent intensity, and SST release. A one sample t test was used for change in membrane potential, change in rheobase, and change in action potential threshold from baseline. An unpaired t test was used to compare membrane potential and changes in membrane potential, rheobase, and action potential threshold between groups. Bonferroni post-hoc tests were used when appropriate. Unpaired t tests and Welch’s t tests were used where appropriate to compare control and binge drinking resting membrane potential, rheobase, and action potential threshold. Data are presented as the mean and standard error of the mean. Total alcohol consumption (g/kg) across all 4 cycles of DID was correlated with SST cell density, immunofluorescent intensity, and change in membrane potential following binge drinking (cells were averaged to create one datapoint per mouse). Control mice were included in correlations with g/kg consumed = 0. Each correlation was reported as Pearson’s r. One outlier was excluded from SST optogenetic release analysis identified by Grubb’s test in Graphpad Prism. Excluding this datapoint did not affect the statistical result. To test for the effects of sex and alcohol consumption on cell density and SSTR+ prevalence in RNAscope experiments, we used ANOVA on independent mixed linear models with fixed effects of sex (2 levels: M, F) and group (2 levels: ETOH, H2O). We regressed fixed effects sex and total ETOH intake on dependent variable SSTR+ transcript density to identify dose-dependent effects of alcohol consumption on transcript density. Across all mixed linear models, region and animal ID were used as random effects, and Sattherwaite’s method was used to estimate degrees of freedom in t- and F-tests.

## RESULTS

### Binge-like alcohol consumption does not alter SST cell density throughout the PFC

We first confirmed in our hands that the DID paradigm produces the typically seen behavioral adaptations related to PFC function 24 hr following cessation of alcohol consumption (**Supplementary Figures 1 and 2**). C57BL/6J male and female mice were subjected to four weeks of DID or control conditions, and mice showed expected relevant deficits in exploratory behaviors (**Supplementary Figures 1 and 2**).

Then, because previous work suggests SST cell density is sensitive to stress paradigms [32–34], we interrogated the effect of binge alcohol consumption on SST cell number in the anterior cingulate cortex (ACC), PL cortex, and IL cortex (**Figure 1A**). 24 hr following binge-drinking, mice were perfused and brains were sectioned, imaged, and cell density quantified (representative images from SST-Cre-Ai9 mice, male and female, control and binge drinking, in **Figure 1B**). There was no significant effect of sex or binge drinking on SST cell density in the ACC (2-way ANOVA; F_sex (1,14)_ = 2.494, *p* = 0.1366; F_binge drinking(1,14)_ = 4.095e-005, *p* = 0.9950, F_sex x binge drinking(1,14)_ = 0.3977, *p* = 0.5375; **Figure 1C**). Total alcohol consumption across the DID paradigm was not significantly correlated with SST cell density in the ACC in males (r(8) = 0.107, *p* = 0.4291) or females (r(10) = 0.0507, *p* = 0.5319; **Figure 1D**) or with the final 4 hr binge in males (r(8) = 0.145, *p* = 0.3525) or females (r(10) = 0.107, *p* = 0.3573; **Figure 1E**). There was no significant effect of sex or binge drinking on SST cell density in the PL (2-way ANOVA; F_sex (1,14)_ = 2.329, *p* = 0.1493; F_binge drinking(1,14)_ = 0.2287, *p* = 0.6399, F_sex x binge drinking(1,14)_ = 0.1587, *p* = 0.6963; **Figure 1F**). Total alcohol consumption across the DID paradigm was not significantly correlated with SST cell density in the PL in males (r(8) = 0.00588, *p* = 0.8567) or females (r(10) = 0.0771, *p* = 0.4372; **Figure 1G**), or with the final 4 hr binge in males (r(8) = 0.00306, *p* = 0.8964) or females (r(10) = 0.126, *p* = 0.3144; **Figure 1H**). There was no significant effect of sex or binge drinking on SST cell density in the IL (2-way ANOVA; F_sex (1,14)_ = 0.1068, *p* = 0.7486; F_binge drinking(1,14)_ = 0.2469, *p* = 0.6270, F_sex x binge drinking(1,14)_ = 0.01028, p = 0.9207; **Figure 1I**). Total alcohol consumption across the DID paradigm was not significantly correlated with SST cell density in the IL in males (r(8) = 0.0263, *p* = 0.7012) or females (r(10) = 0.00711, *p* = 0.8169; **Figure 1J**), or with the finale 4 hr binge in males (r(8) = 0.0206, *p* = 0.7344) or females (r(10) = 0.0229, *p* = 0.6767; **Figure 1K**). These results suggest that SST cell density is unchanged in the medial PFC broadly (ACC, PL, and IL subregions) following binge drinking, highlighting a key difference between stress-induced changes [34] and alcohol-induced changes in PFC cell density. This finding is also consistent with studies in rats which indicate no change in total neuron number (not specific to SST) following a similar model of binge drinking [35].

**Figure 1.**
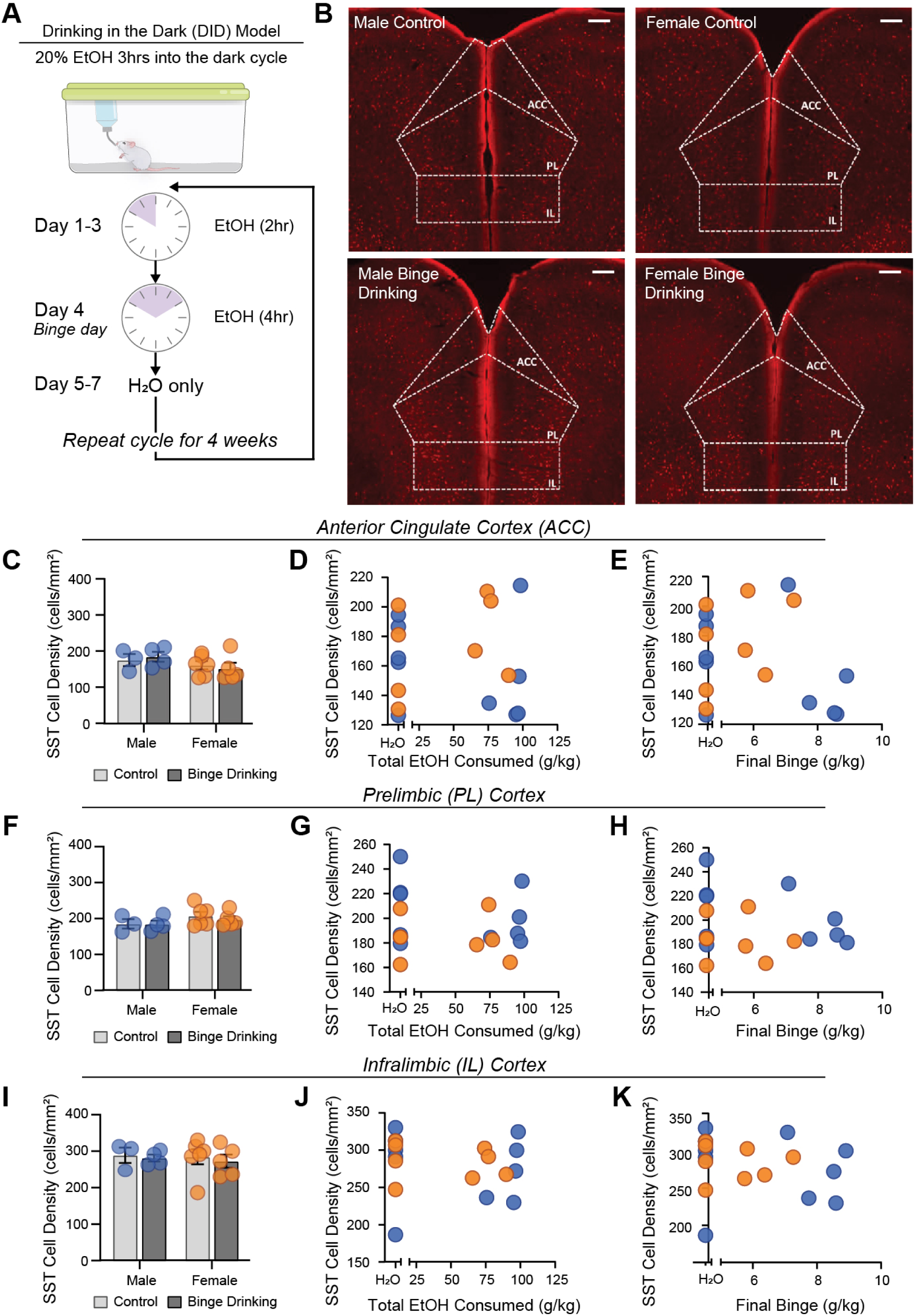
Binge alcohol consumption via the drinking in the dark (DID) model does not alter SST cell density across the mPFC in either male or female mice. (A) Overall timeline and (B) representative images from the ACC, PL, and IL cortex of male and female, control and binge drinking SST-Ai9 mice. Within the ACC, there was no relationship between SST cell density and alcohol consumption (C) regardless of sex, total EtOH consumed (D) or EtOH consumed at the final binge (E). Within the PL cortex, there was no relationship between SST cell density and alcohol consumption (F) regardless of sex, total EtOH consumed (G) or EtOH consumed at the final binge (H). Within the IL cortex, there was no relationship between SST cell density and alcohol consumption (I) regardless of total EtOH consumed (J) or EtOH consumed at the final binge (K). Thus, voluntary alcohol consumption during adulthood does not reduce the overall number of PFC SST neurons, contrasting known stress-induced reductions in cortical SST cell density.

### Binge drinking abolishes SST peptide effects in the PL, but not IL, subregion of the PFC

We next sought to interrogate the effect of binge alcohol consumption on SST peptide modulation of pyramidal neuron function in both the PL and IL cortices. Male and female C57/BL/6J mice underwent four weeks of DID or control conditions, and 24 hr after the final binge session mice were sacrificed for electrophysiology (**Figure 2A**). Pyramidal cells in the PL cortex were identified by their morphology (triangular soma with an apical dendrite) and membrane properties [4, 6, 18]. There was no significant difference in the resting membrane potential (RMP) at baseline (prior to SST application) between control and binge drinking groups (unpaired t test; t_28_ = 0.6974, *p* = 0.4913; **Figure 2B**) consistent with our previously published work [6]; however, the binge drinking exposed cells exhibited larger variability in resting membrane potential (standard deviation; 10.73 mV) than control cells (standard deviation; 4.196 mV) suggesting some (potentially output-pathway dependent) dysregulation in basal excitability following binge drinking (F_(13,15)_ = 6.544, *p* = 0.0009). SST bath application significantly hyperpolarized pyramidal cells in control mice, but this effect was not observed in binge drinking mice (one-sample t test; control t_15_ = 4.665, *p* = 0.0003; binge drinking t_13_ = 1.140, *p* = 0.2747; **Figure 2C-D**). No significant sex differences were identified, with no difference in RMP regardless of drinking condition (two-way ANOVA, F_sex(1,23)_ = 0.6708, *p* = 0.4212; F_binge drinking(1,23)_ = 0.1371, *p* = 0.7146; **Figure 2E**). There was also no correlation between RMP and total alcohol consumption (females r(15) = -0.3675, *p* = 0.146; males r(11) = 0.03198, *p* = 0.9174; **Figure 2F**) or alcohol consumption at last binge (female r(15) = - 0.1939, *p* = 0.4559; male r(11) = -0.05285, *p* = 0.8639; **Figure 2G**) in either sex. Furthermore, the change in RMP induced by SST was similar in both male and female mice (one-sample t test; male control t_6_ = 3.333, *p* = 0.0157; females control t_8_ = 3.794, *p* = 0.0053; males binge drinking t_5_ = 1.290, *p* = 0.2536; female binge drinking t_7_ = 0.2374, *p* = 0.8191; **Figure 2H**). Significant correlations emerged in female mice, with total alcohol consumption across the four-week period correlating with a significantly reduced SST-induced hyperpolarization (females r(15) = 0.5552, *p* = 0.0207; **Figure 2I**), but this effect did not hold true for males (males r(11) = -0.1228, *p* = 0.6894; **Figure 2I**). Alcohol consumed at the last binge did not correlate with SST-induced hyperpolarization in either sex (females r(15) = 0.2060, *p* = 0.4276, males r(11) = -0.07190, *p* = 0.8154 **Figure 2J**). Together, this points towards alcohol-induced dysregulation of SST peptide efficacy in the PL cortex of male and female mice, in a manner that may be more clearly dose-dependent in females.

**Figure 2.**
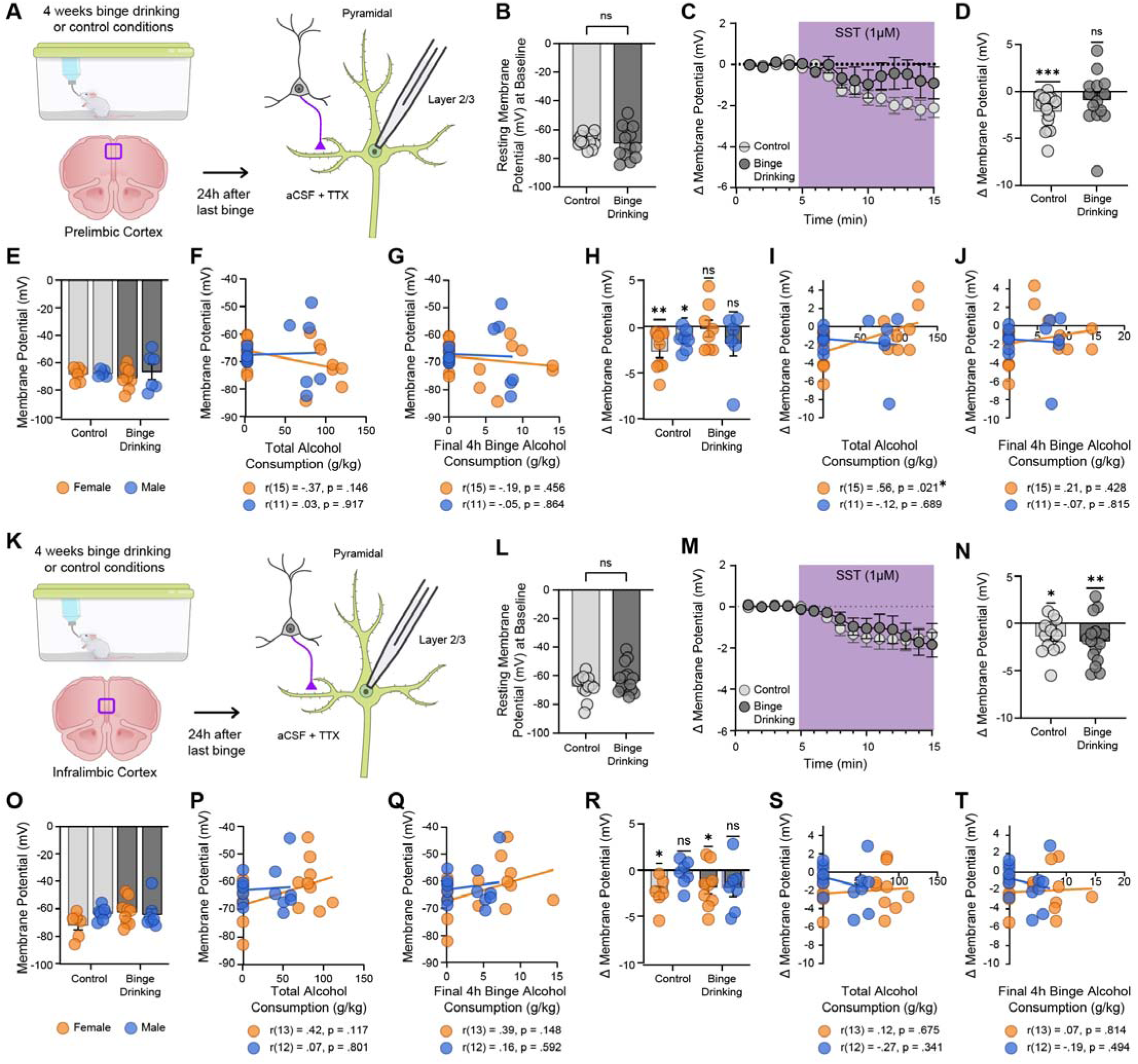
Binge alcohol consumption reduces SST-induced pyramidal neuron hyperpolarization in the PL, but not IL, cortex in both male and female mice, and is driven by total alcohol consumption in female mice. (A) Experimental procedure. Adult male and female mice underwent four weeks of DID and PL electrophysiology was performed 24 hr after the last binge exposure. TTX was added to the aCSF to block action potentials and polysynaptic effects. Cells were held in current clamp configuration and the resting membrane potential (RMP) was recorded. (B) There was no significant difference in PL pyramidal neuron RMP between control and binge drinking groups. (C) Change in membrane potential across time following 1 μM SST bath application. (D) 1 μM SST significantly hyperpolarizes pyramidal cells from control mice but not binge drinking mice. (E). There was no correlation between the PL RMP and total alcohol consumed (F) or alcohol consumed at the final binge (G) in either sex. The deficits in 1μM SST-induced hyperpolarization occurred regardless of sex (H). Interestingly, the change in RMP induced by DID correlated with overall alcohol consumed in females but not males (I). Final binge did not correlate with alcohol consumed (J). Identical experiments were conducted in the IL cortex (K). There was no significant difference in IL pyramidal neuron RMP between control and binge drinking groups (L). (M) Change in membrane potential across time following 1μM SST bath application. (N) 1 μM SST significantly hyperpolarized pyramidal cells from both control mice and binge drinking mice. When assessing sex differences, there was no significant difference in IL pyramidal neuron RMP across drinking groups and sex (O). There was no correlation between the IL RMP and total alcohol consumed (P) or alcohol consumed at the final binge (Q) in either sex. However, interesting sex differences emerged when accounting for the SST-induced change in RMP in the IL cortex. While both control and binge drinking female mice showed hyperpolarization following 1 μM SST bath application, this peptidergic effect did not occur in male mice regardless of alcohol condition, suggesting sexual dimorphism in the IL cortex SST system (R). There was no correlation between the change in RMP driven by SST and total alcohol consumption in either sex (S). There was also no correlation between the change in RMP driven by SST and alcohol consumed at the

Identical experiments were performed on pyramidal cells in the IL cortex (**Figure 2K**). Similar to the PL cortex, there was no significant difference in the RMP at baseline between control and binge drinking groups (unpaired t test; t_27_ = 1.245, *p* = 0.2237, **Figure 2L**). SST significantly hyperpolarized pyramidal cells in the IL cortex in both control and binge drinking groups (one-sample t test; control t_12_ = 2.598, *p* = 0.0233; binge drinking t_15_ = 3.068, *p* = 0.0078; **Figure 2M-N**) suggesting that SST signaling in the IL cortex remains intact following binge-like alcohol consumption. There was no significant difference in the RMP of pyramidal regardless of drinking condition or sex (two-way ANOVA, F_sex(1,26)_ = 0.6661, *p* = 0.4218; F_binge drinking(1, 26)_ = 1.725, *p* = 0.2005; **Figure 2O**), and this did not correlate with total alcohol consumed across the four-week period (females r(13) = 0.4217, *p* = 0.1174; males r(12) = 0.07439, *p* = 0.8005; **Figure 2P**), nor with alcohol consumed during the final binge (females r(13) = 0.3926, *p* = 0.1477; males r(12) = 0.1572, *p* = 0.5916; **Figure 2Q**).

Surprisingly, sex differences were revealed in the SST-driven control over IL pyramidal neurons. When data was split by sex, female mice showed a robust SST-driven hyperpolarization that was intact following binge drinking (females control t_5_ = 3.487, *p* = 0.0175; females binge drinking t_8_ = 2.346, *p* = 0.0470; **Figure 2R**). Male mice, however, showed no SST-mediated response at baseline, and this lack-of-response remains following alcohol consumption (males control t_6_ = 0.6787, *p* = 0.5226; males binge drinking t_6_ = 1.843, *p* = 0.1148; **Figure 2R**). SST-driven changes in RMP in the IL cortex did not correlate with total alcohol consumed (females r(13) = 0.1180, *p* = 0.6754; males r(12) = -0.2754, *p* = 0.3405; **Figure 2S**) or alcohol consumed at last binge (females r(13) = 0.06643, *p* = 0.8140; males r(12) = -0.1995, *p* = 0.4940; **Figure 2T**) in either sex. This suggests that while alcohol does not appear to interfere with SST signaling in the IL cortex, there are robust basal sex differences in this system that warrant further investigation. Specifically, female circuits appear to engage SST peptide-mediated control over PFC outputs, whereas males do not.

### Binge alcohol consumption does not disrupt abundance or evoked release of SST peptide in the PFC

Given the dysregulation of SST signaling in the PL cortex following binge drinking, we next sought to elucidate the role of locally released SST in the PL cortex on pyramidal cells. We first examined whether overall SST peptide abundance was altered in the PL cortex (representative images in **Figure 3A**). In the PL, alcohol consumption did not induce overall changes in SST immunofluorescence intensity in either sex (2-way ANOVA F_sex(1,21)_ = 0.4422, *p* = 0.5133; F_binge drinking (1,21)_ = 0.4533, *p* = 0.5081; **Figure 1B**) and was not influenced by total ethanol consumed in females (r(10) = 0.110, *p* = 0.0727; **Figure 3C**) or males (r(10) = -0.507, *p* = 0.0924; **Figure 3C**). Similarly, no correlation was seen with ethanol consumed at final binge in females (r(10) = 0.0908, *p* = 0.07789; **Figure 3D**) or males (r(10) = -0.539, *p* = 0.0704; **Figure 3D**).

**Figure 3.**
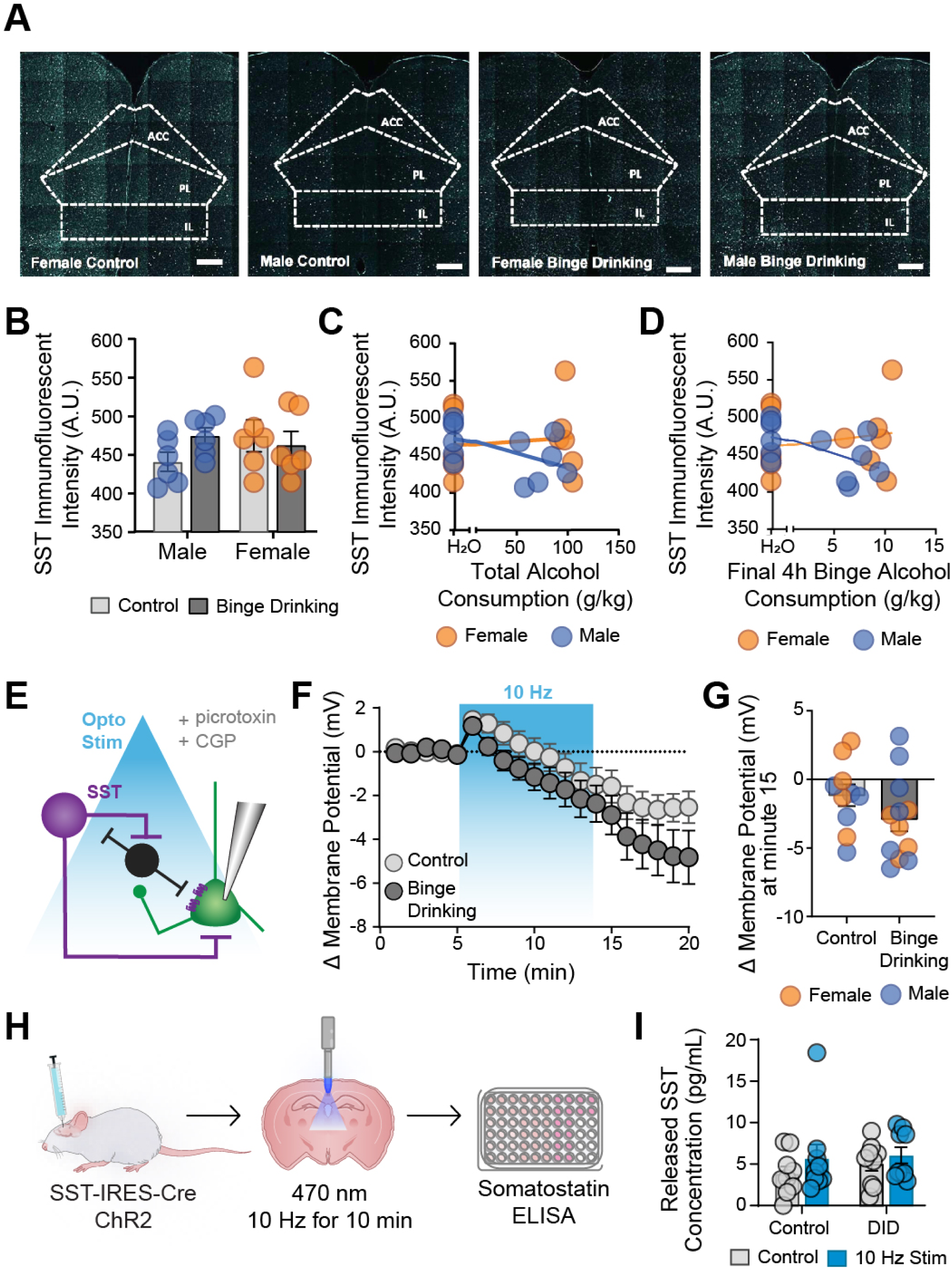
Binge drinking does not alter SST peptide abundance, or the capacity for SST release, in the PL cortex of either male or female mice. (A) Representative images of SST IHC from male and control, female and binge drinking mice. (B) There was no effect of binge drinking on PL SST expression regardless of sex. There was no correlation between SST expression and total alcohol consumed (C) or alcohol consumed at last binge (D) regardless of sex. (E) Overview of optogenetic stimulation slice electrophysiology experiments. Picrotoxin and CGP were included in the aCSF to block GABA receptor mediated effects. Change in PL pyramidal RMP across time following optogenetic activation (10 Hz) of SST neurons (F). There was a significant optogenetically-induced hyperpolarization in both control and binge drinking mice (G) with no significant sex differences. (I) overview of optogenetic stimulation ELISA experiments. (J) There was no significant difference in baseline or optogenetically-evoked SST levels regardless of alcohol condition.

We next examined the impact of optogenetically-evoked endogenous SST release on the RMP of pyramidal cells in both control and binge drinking mice by employing prolonged optogenetic stimulation (10 Hz for 10 minutes). We isolated peptidergic contributions by inhibiting local GABAergic receptor actions (though other potentially co-released peptides were not blocked). 24 hr following the last binge, SST-Ai32 mice (expressing channelrhodopsin in SST cells) were prepared for electrophysiology (**Figure 3E**). Layer 2/3 pyramidal neurons were patched, and following a 5 min stable baseline, SST cells were optogenetically stimulated while RMP was continuously recorded (**Figure 3E**). Prolonged optogenetic activation of SST cells resulted in no significant difference between control and binge drinking groups (unpaired t test; t_20_ = 1.437, p = 0.1661; **Figure 3G**). Variances did not differ between control and alcohol-exposed groups (p = 0.550). There were no differences seen across sexes. Using a complementary plate-based ELISA (enzyme-linked immunosorbent assay; **Figure 3H-I**), optogenetic-induced SST release into the aCSF was not altered following binge drinking (two-way ANOVA, F_alcohol (1,32)_ = 0.4777, *p* = 0.4944; F_opto stimulation (1,32)_ = 1.580, *p* = 0.2179). Together, these experiments suggest that binge drinking does not reduce the *capacity* for SST peptide release in the PL cortex, further suggesting post-synaptic mechanisms underlying microcircuit changes.

### Deficits in PL cortical SST functioning following DID are driven in part by a loss of SSTR4 expression

Our experiments thus far suggested a profound dysregulation in the PL SST system that was not driven by changes in abundance or potential release of SST peptide. We therefore explored if changes in SST function were driven by alterations of expression in SST receptors. While SST has five known endogenous receptors, our experiments targeted SSTR4 due to its abundance in the cortex and accessibility for assessment via RNAscope (see **Supplementary Figure 3 and 4** for image processing pipeline). First, we tested whether binge drinking decreased cell counts of Sstr4+ cells in the PL cortex. Exposure to binge drinking did not affect cell counts (F_(1,12.03)_ = 0.037, *p* = 0.85) or prevalence of Sstr4+ cells (F_(1,11.9)_ = 0.031, *p* = 0.58; **Figure 4A**), complementing results reported in Figures 1 and 3. We then asked whether *Sstr4* transcription was altered following alcohol consumption. In Sstr4+ cells, we found that alcohol significantly reduced Sstr4 transcript density in female animals, but not male animals (t_39_ = 2.467, *p* = 0.018; **Figure 4B**). In females, greater ethanol consumption led to greater reductions in Sstr4 transcript density, whereas males exhibited no dose-dependent modulation (females: t_39_ = -2.92, *p* = 0.005; **Figure 4D**). This suggests that the reductions in SST-induced hyperpolarization seen following binge drinking are driven by reduced transcription of *Sstr4* in female mice, and complement the consumption-dependent effects see in females in Figure 2.

**Fig 4.**
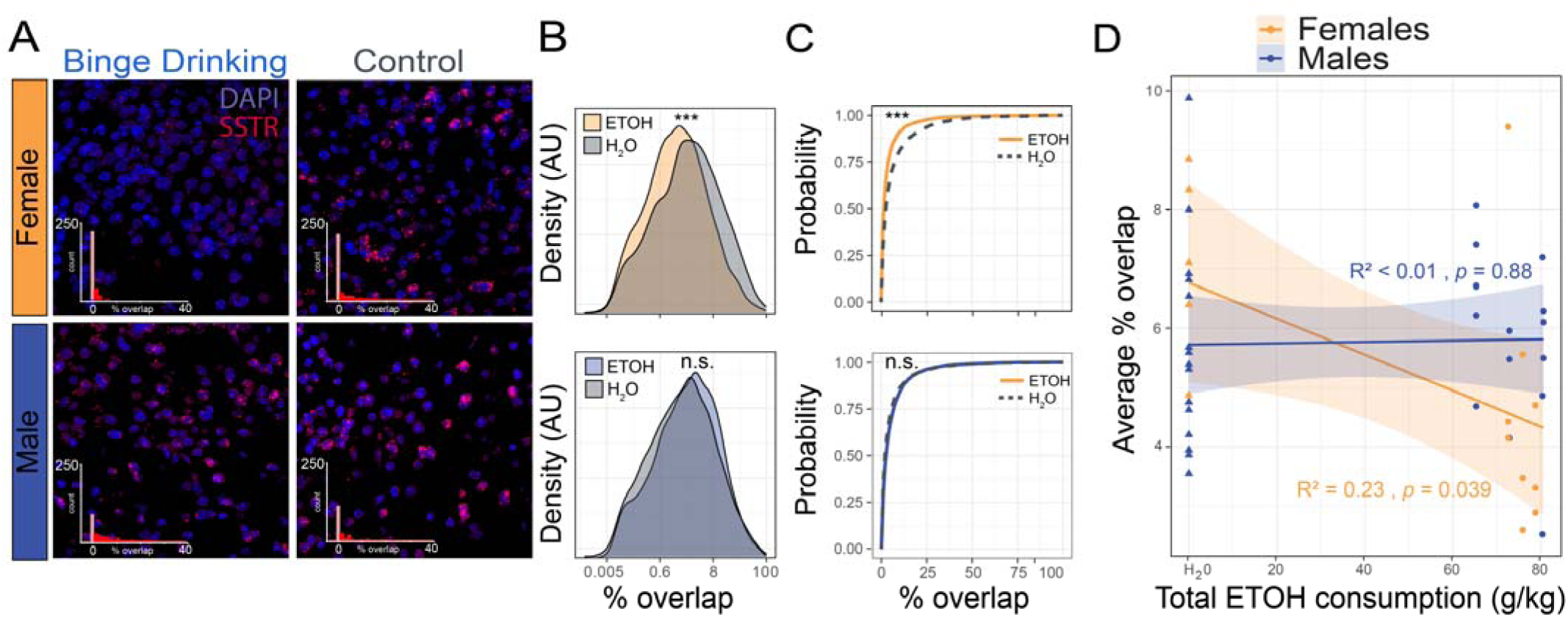
Binge drinking reduces SSTR-encoding transcript density in female mice. (A) Representative images of male and female mice from control and DID conditions. Inset histograms count DAPI regions binned by percent overlap with SSTR. (B) Estimated probability densities of percent overlap in SSTR+ DAPI regions. (C) Empirical cumulative distribution functions for percent overlap across all DAPI regions. (D) Regression of total ETOH consumption on average % overlap across all

## DISCUSSION

Our current work comprehensively characterizes how a well-validated animal model of binge drinking (drinking in the dark, DID) alters SST peptidergic control over cortical circuits (**Figure 5**). 24 hr following four weeks of DID, pathway-agnostic assessment of pyramidal output neurons revealed those in the PL, but not IL, cortex had diminished responsiveness to SST peptide bath application, as measured by changes in membrane potential. This is not driven by reduced number of SST positive neurons (as seen with stress), clear deficits of SST peptide abundance or capacity for peptide release but instead appears to be driven in females by a genetic downregulation of SSTR4. While a clear relationship did not emerge in males, deficits in females scaled with alcohol dose. This suggests that pharmacological targeting of the SST system to restore post-drinking deficits will require restoration of receptor levels, potentially through pharmacological mechanisms targeting the GPCR signaling cascades [36]. While SSTR4 is abundantly expressed in the cortex, other subtypes are as well [37] and future work will investigate the contribution of other SSTR subtypes.

**Fig. 5.**
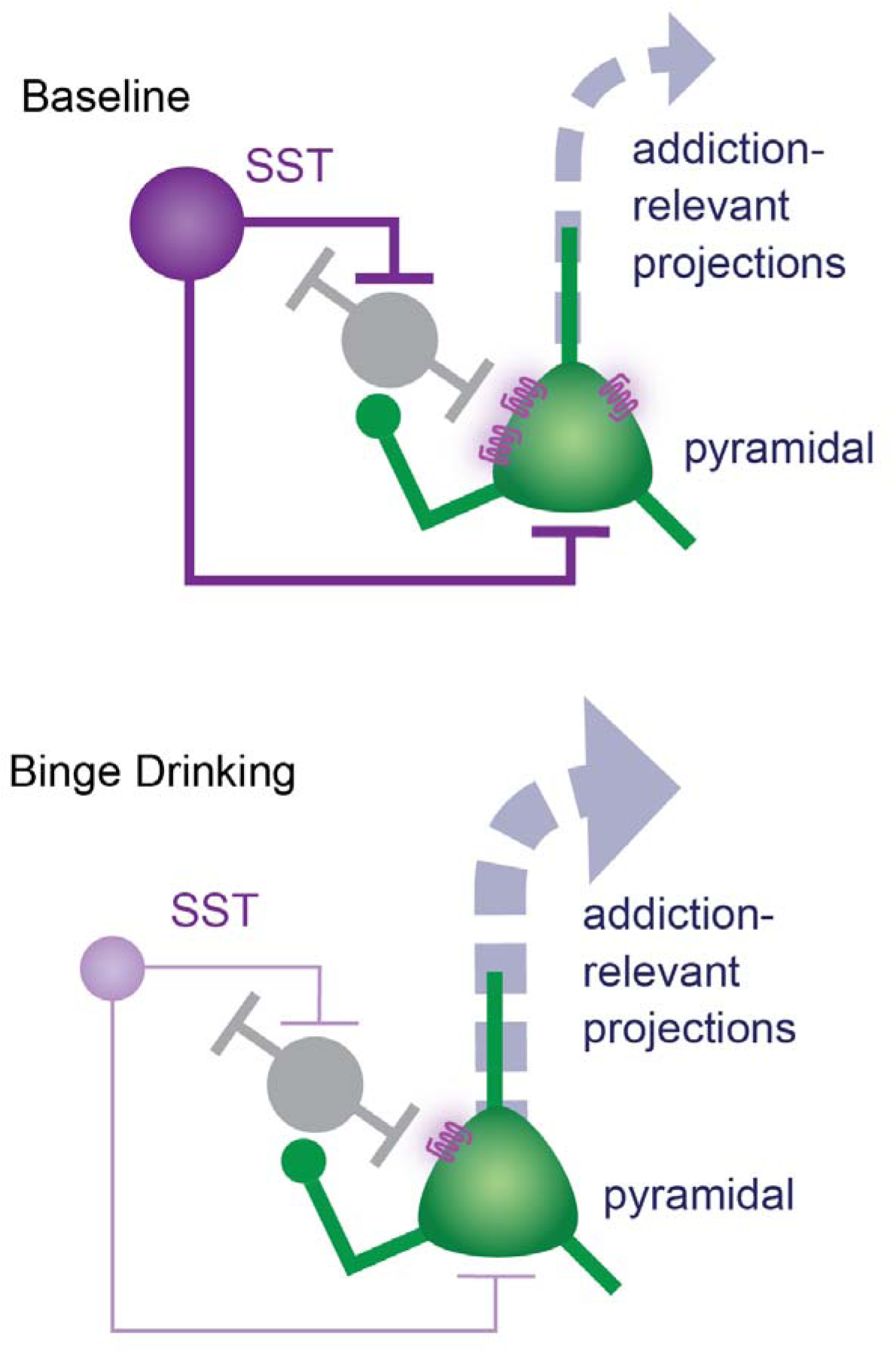
Overall framework for alcohol-induced disruptions in PFC SST peptide control over local neurocircuits. In baseline conditions, PL cortical SST peptide signaling provides robust inhibitory control over PL pyramidal neurons, providing a mechanism to regulate PFC engagement. Following binge drinking, our previous work shows downregulation of SST intrinsic excitability and GABAergic control over pyramidal neurons. Here, we extend this to alcohol-induced downregulation of SST peptidergic control. This sets the stage for PFC pyramidal hyperexcitability, which is one putative pathway underlying the transition from causal substance use to addiction.

SST neurons within the mPFC have the potential to serve as master regulators of PFC function and output activity, particularly for affective states [38], and their peptide signaling promotes exploratory behaviors via a reduction in overall cortical activity in the PFC [4]. Due to their role in controlling multiple behavioral states, their disruption by external stimuli such as drugs of abuse have the potential to profoundly disrupt circuit dynamics and overall behavioral function. Recent work highlights the diversity of SST cells, suggesting distinct morphological and functional categories [39–42]. The function of SST neurons on cortical outputs differs by SST subpopulation [43], and optogenetic activation of large portions of SST cells, as well as bath application of the peptide, likely does not recapitulate the endogenous firing and release dynamics of this diverse population of cells. There is also dense overlap with other neuropeptides such as dynorphin [44]. The development of optical biosensors for neuropeptides [45] will allow deeper investigation into the spatial and temporal topography of neuropeptide actions. Together, our work provides a clear mechanistic deficit in PFC SST peptidergic function following a clinically- and translationally-relevant model of alcohol consumption.

## ACKNOWLEDGEMENTS AND FUNDING

R01AA029403, The Department of Biology, Brain and Behavior Research Foundation NARSAD Young Investigator Award to NAC; R01NS078168 to PJD; R01AG074041 to JLK, T32NS115667 and AHA 24PRE1201066 to MSH, AG049676 to LB, and F32AA031396 to LRS.

## DATA AVAILABILTY STATEMENT

All data and code will be made upon request

## CONTRIBUTIONS

DFB and NAC designed the experiments. DFB, KRG, AK, LB, LRS, and MU conducted the experiments. DFB, KRG, JMR, and MSH contributed to analysis. DFB and NAC wrote the manuscript, with input from all authors. JLK, PJD, and NAC were responsible for funding acquisition.

## Supplementary Methods

### Exploratory behavior testing

24 hours post-DID, a short battery comprised of open field testing (OFT) and elevated plus maze (EPM) assessment was used to investigate changes in exploratory behavior after DID (**Supplementary Figures 1 and 2**). Tests were conducted in the dark phase of the daily light cycle under red light. Behavior was recorded by a camera mounted above the arenas. First, mice were transferred to a procedure room and allowed to acclimate for at least an hour before behavioral testing. Each mouse was placed in the corner of the open field (50 × 50 cm; black plexiglass 20 cm walls) and allowed to freely explore for 5 min. After testing, mice were returned to home cages and were not disturbed for at least one hr. Next, each mouse was placed in the center of an elevated plus maze (40 cm elevation; 30 × 5 cm arms; clear plexiglass 20 cm walls on closed arms) and monitored for 5 min. Behaviors were tracked using DeepLabCut (Brockway et al., 2023) and statistical analysis was conducted in MATLAB. Generalized linear mixed-effect models were used to test the associations between exploratory behaviors with DID treatment assignment and, among alcohol-exposed subjects, total alcohol and last binge alcohol consumed. Behavior was tested in multiple cohorts, which was included as a random effect in the analysis. For assessment of DID treatment effects, the model was: [outcome variable] ∼ 1 + DID*Sex + (1|Cohort). To assess total alcohol and last binge alcohol effects within alcohol-exposed groups, the model was: [outcome variable] ∼ 1 + Total_EtOH*Sex + Last_Binge_EtOH + (1|Cohort).

### Somatostatin immunohistochemistry for cell density

Mice were deeply anesthetized with Avertin (250 mg/kg) and perfused transcardially with ice-cold phosphate buffered saline (PBS, pH 7.4) and 4% paraformaldehyde (PFA, pH 7.4). Brains were removed, post-fixed in PFA overnight washed 3x in PBS and stored in PBS at 4 C for less than 1 week. 40-μm free floating sections containing the PFC were sliced with a vibrating microtome (Leica, VS 1200) and stored in PBS until staining, conducted within one week.

Prior to immunostaining, slices were washed 3 times with PBS for 10 min each on a Fisherbrand™ Multi-Platform Shaker, then underwent antigen retrieval in 10 mM sodium citrate buffer (pH 6.0) at 80°C for 30 min. Slices were washed three times in PBS for 10 min each, and permeabilized in 0.5% Triton X-100 in PBS for 60 min. Nonspecific binding was blocked with 5% normal goat serum (NGS) in 0.1% Triton X-100 in PBS for 60 min. Slices were washed 3 times in PBS for 10 min each, then incubated in a primary antibody rat anti-somatostatin (1:500, Millipore, Burlington, MA, United States) in 2.5% NGS in 0.1% Triton X-100 in PBS for 48-h at 4°C. Slices were washed three 3 times with PBS for 10 min each, and incubated in a fluorophore-tagged secondary antibody goat anti-rat Cy3 (1:500, Millipore, Burlington, MA, United States) for 4-h at room temperature. Slices were washed again three times with PBS, with the last step including DAPI (1:10,000). Slices were mounted on glass slides, air-dried and coverslipped with Immunomount (Thermo Fisher Scientific, Waltham, MA, United States). Images were obtained at 20x with an Olympus BX63 upright microscope (Center Valley, PA, United States) under matched exposure settings. Eight images from the PFC were taken per mouse.

SST+ cell counts were performed using ImageJ (National Institutes of Health, Bethesda, MD, United States). The anterior cingulate cortex (ACC), prelimbic (PL) cortex and infralimbic (IL) cortex were delineated and SST neurons were quantified in each region separately under matched criteria for size, circularity, and intensity consistent with our previously published work (Dao et al., 2020; Suresh Nair et al., 2022). The threshold for delineation of cells was set at 8.10±.01. Each ROI’s total SST cell count was divided by the ROI area to give a total SST density represented in cells/mm^2^. 5-8 sections containing the ACC, PL, and IL were quantified and averaged to obtain one value per mouse. Imaging, cell counting, and analysis was done blinded to condition and sex. Original images were used for quantification and representative images were adjusted for brightness and contrast.

### RNAscope

Fresh-frozen brains were sliced in 20 μm coronal sections between coordinates 1.3 mm to 1.9 mm relative to Bregma to capture the PL cortex. Slides were prepared to include anatomically matched sections from each experimental group on each slide. RNAscope was performed on three slides for each animal, as previously published (Wang et al., 2012) using the RNAscope Multiplex Fluorescent v2 Assay kit. Slides were kept in 100% ethanol overnight at -20°C following the fixation and dehydration steps. Slides were then stained for SSTR4 using the RNAscope Probe-Mm-SSTR4-C2 (ACDBio, #416641-C2), diluted 1:50 in RNAscope Probe Dilutant (ACDBio, # 300041). Slides were counterstained with DAPI (provided in the RNAscope kit). Opal 690 (Akoya Biosciences, #FP1497001KT) was used for the SSTR4 probe. One image was taken from each hemisphere of using a STELLARIS 5 white light laser confocal microscope (Leica) with a 20X objective and 2X zoom. The pinhole, gain, and exposure were consistent for each image on a single slide. Images were taken 1024 x 1024 resolution with 2X line averaging. All staining and imaging were done by an experimenter blinded to the conditions.

### RNAscope image processing

RNAscope images were segmented and measured in ImageJ using a custom Python library (**Supplementary Figure 3**). The semi-automated pipeline required user inspection of each individual image, and selection of a few key parameters to be used in fully automated processing for the second stage. First, each two-channel color confocal image was separated into DAPI and SSTR grayscale channels, respectively. To avoid registration of single pixels, we used a small gaussian blur between 0 and 3 pixels in x and y directions on each image, determined manually by inspection. We found that DAPI image registration improved with gaussian blurs with 2-pixel radius, whereas SSTR images generally required less gaussian blur for accurate registration. These blurred images were then binarized using a threshold function with a range manually selected to maximize retention of visible puncta, while minimizing the effect of spatially varying illuminance from uneven sample thickness or edge artifacts. The binarized DAPI images were processed with a “fill holes” function to close incompletely closed regions, and then a watershed function was used to segment adjoined DAPI regions. Each individual DAPI region was then labeled with an identifier unique to that image. We then saved the selected gaussian blur, watershed, and threshold parameters for each channel, in each image, and used a fully automated algorithm to complete segmentation and ROI measurement of all images using an ImageJ script. This algorithm recorded the area of each individual DAPI region and the area of the overlapping SSTR puncta, then saved these results in a tabulated comma-delimited file, along with accompanying file-specific metadata. All comma-delimited files were then opened, joined and analyzed in R using the ESS package in GNU-Emacs.

After unblinding to animal sex and alcohol condition, we asked whether our manually selected image parameters were randomly biased across these factors by chance. To test if manually-selected gaussian blur values or threshold values varied with animal sex, or alcohol condition, we performed 2-way ANOVA with main effects Sex and Group and random effects Animal ID on these dependent variables. We found that gaussian blur radii in DAPI images were no different between sexes or experimental group (Sex, F[1,11.2]=1.0, p=0.34; Group, F[1,11.2]=0.04 p=0.82). Likewise, gaussian blur parameters selected for SSTR images were the same between mouse sexes and groups (Sex, F[1,10.9]=0.45, p=0.51; Group, F[1,10.9]=0.09, p=0.75). We also tested whether manually-selected threshold values were different between mouse sexes or treatment arms in both DAPI and SSTR images. In DAPI images, the manually selected threshold values did not differ between animal sex or treatment arm (Sex, F[1,532]=0.12, p=0.72; Group, F[1,532]=0.11, p=0.74). The same was true for threshold values in SSTR images (Sex, F[1,11.9]=1.01, p=0.33; Group, F[1,11.9]=0.24, p=0.62; **Supplementary Figure 4**). Thus, manually selected image processing parameters were not different between images from male or female mice, nor between images of ETOH+ and control mice.

We then quantified DAPI registration errors across images. Registration error occurs when regions of extreme size are incorrectly identified as a DAPI region, caused at the extremes by single-pixel registrations or registration of large, improperly segmented regions. To identify registration errors, we log-transformed the experiment-wise distribution of DAPI region area to approximate normality, then defined outliers as DAPI regions with area more extreme than 1.5 IQR of the median (**Supplementary Figure 4C-D**). Using this inclusion criterion, we found no difference in the outlier rate based on animal sex (F[1,11.0]=2.29, p=0.15) or treatment arm (F[1,11.0]=1.88, p=0.19). Nearly all excluded regions were below the minimum acceptable DAPI size (<3.68 LJm2), consistent with the left-skewedness of the log-transformed distribution. This distribution is expected after segmenting DAPI images with a watershed function, as the watershed disproportionately splits larger DAPI regions into smaller ones, skewing the distribution left.

### Patch-clamp slice electrophysiology

Brains were immediately placed in ice-cold oxygenated NMDG-HEPES artificial cerebrospinal fluid (aCSF) containing the following, in mM: 92 NMDG, 2.5 KCl, 1.25 NaH2PO4, 30 NaHCO3, 20 HEPES, 25 glucose, 2 thiourea, 5 Na-ascorbate, 3 Na-pyruvate, 0.5 CaCl2·2H2O, and 10 MgSO4·7H2O (pH to 7.3–7.4). The PL was identified according to the Allen Mouse Brain Atlas. 300 µm coronal slices containing the PL were prepared on a Compresstome Vibrating Microtome VF-300-0Z (Precisionary Instruments, Greenville, NC), and transferred to heated (31°C) NMDG-HEPES (in mM: 124 NaCl, 4.4 KCl, 2 CaCl2, 1.2 MgSO4, 1 NaH2PO4, 10.0 glucose, and 26.0 NaHCO3, pH 7.4, mOsm 300-310), for a maximum of 10 min. Slices were then transferred to heated (31°C) oxygenated normal aCSF where they were allowed to rest for at least 1 hr before use. Finally, slices were moved to a submerged recording chamber where they were continuously perfused with the recording aCSF (2 mL per min flow rate, 31°C). Recording electrodes (3–6 MΩ) were pulled from thin-walled borosilicate glass capillaries with a Narishige PC-100 Puller.

Following rupture of the cell membrane, cells were held in current-clamp. A minimum of 5 min stable baseline was acquired prior to experiments and bath application. Measurements of intrinsic excitability were conducted at both resting membrane potential (RMP) and at the standard holding potential of –70 mV both before and after application. Gap-free RMP was recorded during the entire drug application period. For optogenetic activation of SST cells with simultaneous electrophysiological recording of pyramidal cells in layer 2/3 of the PL cortex, slices from SST-IRES-Cre:Ai32 mice (expressing Cre-dependent channelrhodopsin in SST cells) were kept shielded from light, and experiments performed under low illumination. 25 μM Picrotoxin (Hello Bio, HB0506), and 1 μM CGP 55845 (Tocris, 1248) were added to the aCSF to block GABAergic signaling. Cells were held in current clamp and membrane potential (mV) was measured. Following establishment of a 5 min stable baseline, a 470-nm LED (CoolLED, United Kingdom) was directed to the slice for 10 min (10 Hz frequency).

### Enzyme immunoassay for photostimulated release of SST

24 hr after DID, SST-IRES-Cre:Ai32 mice (expressing Cre-dependent channelrhodopsin in SST cells) were sacrificed for enzyme immunoassay for photostimulated release of SST. Briefly, mice were rapidly decapitated under isofluorance anesthesia and the dissected brain was placed in ice-cold modified high-sucrose artificial cerebrospinal fluid (in mM: 194 sucrose, 20 NaCl, 4.4 KCl, 2 CaCl2, 1 MgCl2, 1.2 NaH2PO4, 10.0 glucose, and 26.0 NaHCO3, pH 7.4). 150 mM coronal slices containing the PL were sectioned on a Compresstome (Precisionary Instruments, Greenville, NC) and transferred to normal aCSF (in mM: 124 NaCl, 4.4 KCl, 2 CaCl2, 1.2 MgSO4, 1 NaH2PO4, 10.0 glucose, and 26.0 NaHCO3, pH 7.4) for 30 min. Slices were maintained in a holding chamber at 31 °C, continuously bubbled with a 95% O2/ 5% CO2 mixture and shielded from light until photostimulation. Individual slices were placed in a well on a 12-well plate with 500 mL of oxygenated normal aCSF. New aCSF was used at this step to control for any SST leakage following slicing. A 470-nm laser (CoolLED, United Kingdom) was directed to the PL cortex for 15 min at a stimulation frequency of 10 Hz. Stimulation/control conditions were counterbalanced between rostral and caudal PL slices to account for variability in viral expression along the anterior/posterior axis. No peptidase enzyme inhibitor was added to the surrounding aCSF. Samples were kept on ice until processed for enzyme linked immunoassay (ELISA).

At the end of the experiment, 2 x 50 mL of the aCSF within the well was pipetted into a 96-well SST ELISA plate (LSBio Somatostatin ELISA Kit LS-F12622) and ELISA (including standard curve) were performed identical to manufactures instructions. The assay is specific for mature forms of SST, somatostatin-14 and somatostatin-28, with no cross-reaction to similar peptides such as neuropeptide Y and vasoactive intestinal peptide. Absorbance was read at 450 nm on a Synergy2 microplate reader (BioTek, Winooski, VT). SST release was quantified in pg/mL. Following optical stimulation, slices were visualized under infrared video microscope and blue LED (470 nm) for verification of viability and eYFP expression in the PL. Photostimulation, ELISA, and analysis were performed blinded to condition.

## Supplementary Figures

**Supplementary Figure 1.**
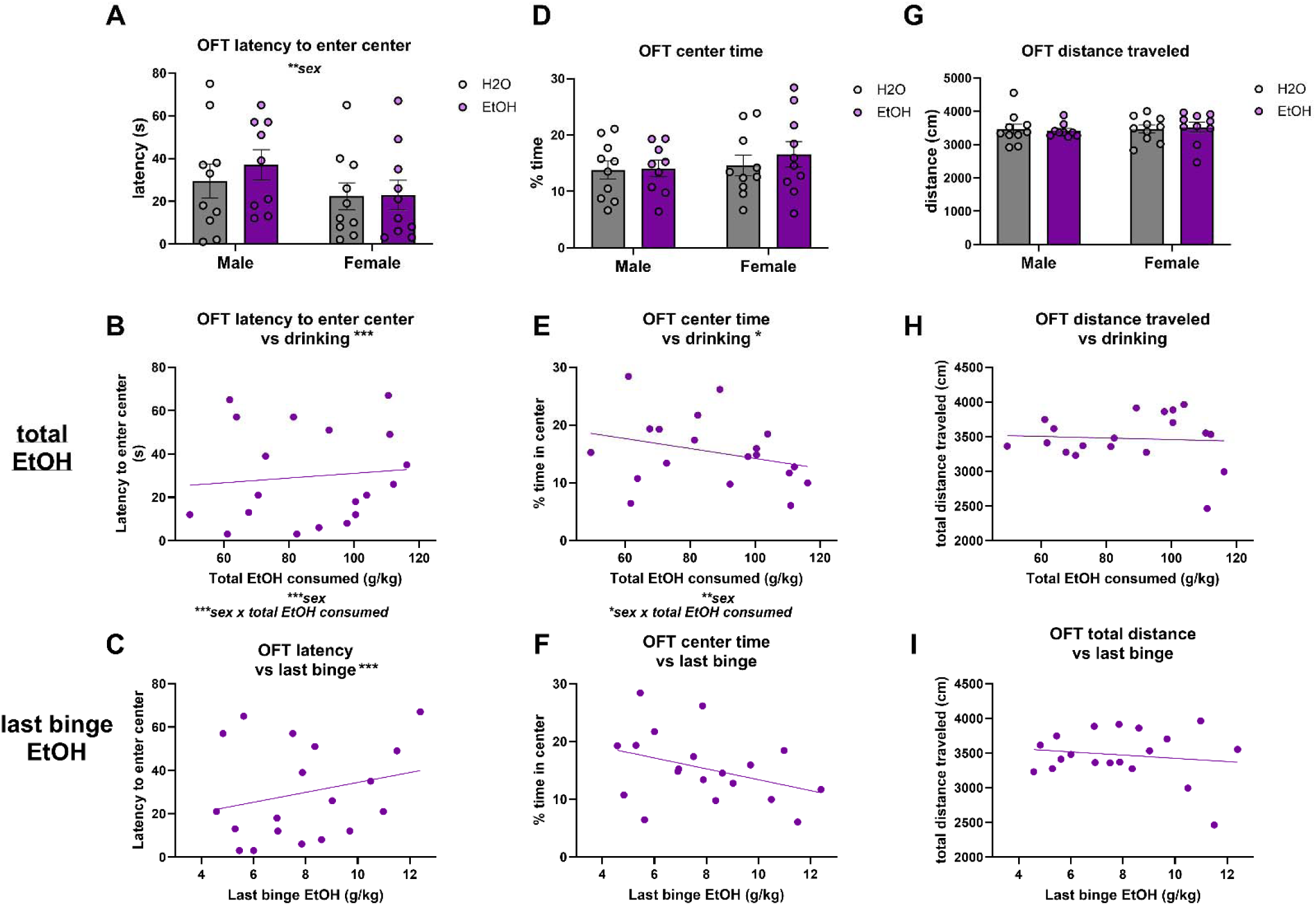
Exploratory behaviors in the OFT 24 hours post-DID. Among DID mice, total and last binge alcohol consumption were inversely associated with exploratory behaviors 24 hr post-DID. (A) Across all conditions, latency to enter the center was higher in males than females (male: 0.2798 *±* 0.088736, p = 0.003). (B-C) Total alcohol consumed (0.05166 *±* 0.0095662, p<0.001) and last binge alcohol consumed (0.16692 *±* 0.036066, p<0.001) were positively associated with latency to enter center. Again, males exhibited a longer latency to enter the center than females (male: 7.0737 *±* 0.99888, p<0.001). An interaction of sex and total alcohol consumed was also detected (male x total EtOH: −0.059403 *±* 0.0097241, p<0.001). (D) No effects of DID or sex were detected. (E-F) Total alcohol consumed was inversely associated with time spent in the center (−0.30748 *±* 0.1125, p=0.016), which was also influenced by an interaction of sex and total alcohol consumed (male x total EtOH: 0.3369 *±* 0.12171, p=0.015). Independent of alcohol effects, males spent less time in the center than females (male: −36.529 *±* 10.263, p = 0.003). (G-I) Total distance traveled in the open field was not found to be influenced by sex or alcohol. For scatterplots, lines of best fit were added independently of the described analysis to aid visualization. * = p<0.05, ** = p<0.01, *** = p<0.001

**Supplementary Figure 2.**
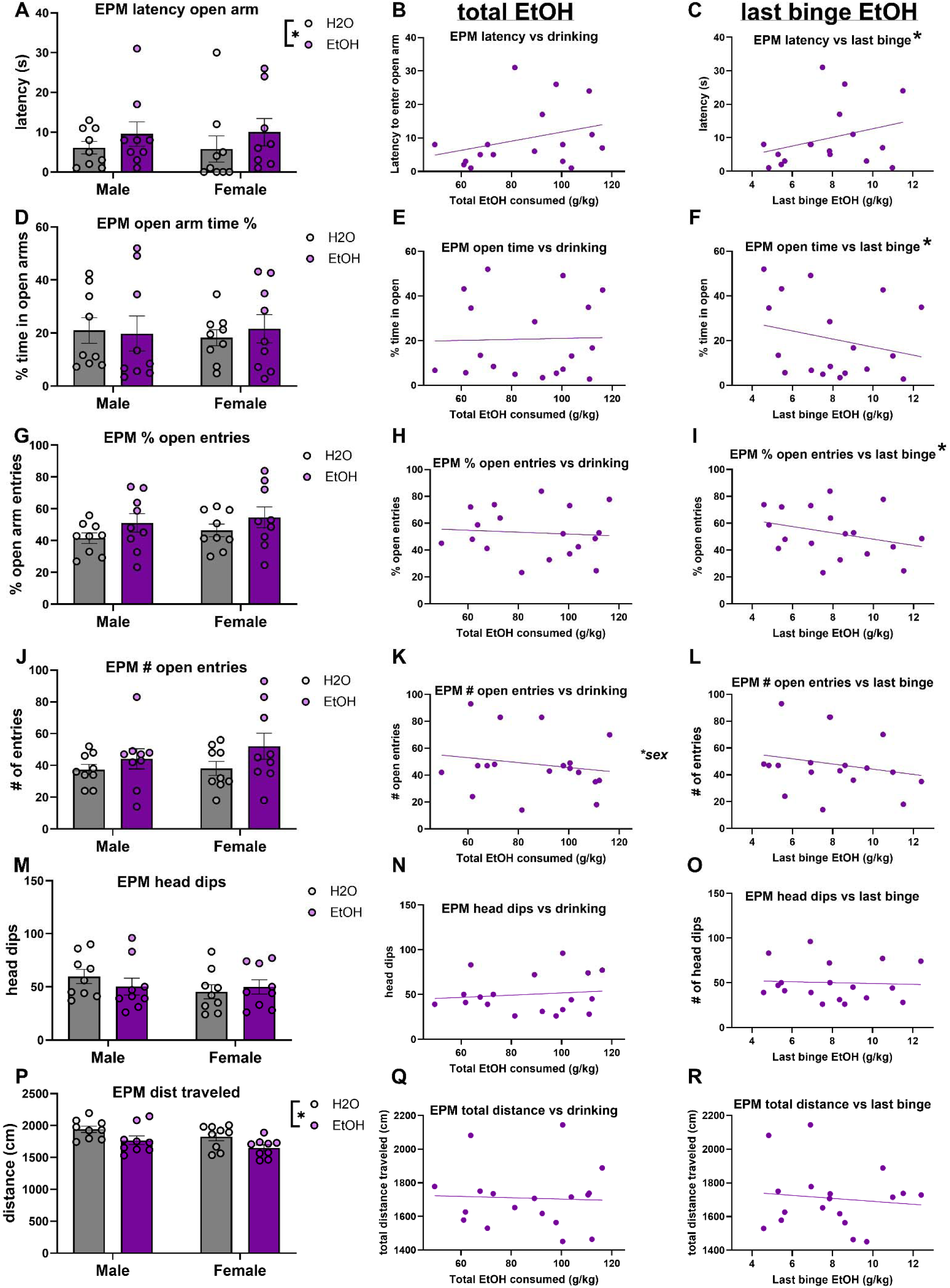
Exploratory behaviors in the EPM 24 hours post-DID. Alcohol exposure reduced exploratory behaviors in the elevated plus maze 24 hr post-DID, and last binge alcohol consumption strongly predicted decreases in exploration. (A-C) Mice that consumed alcohol had an increased latency to enter an open arm relative to water-exposed control subjects (EtOH: 0.50305 *±* 0.17824, p=0.008). Among alcohol-exposed subjects, alcohol consumed during the last binge was positively associated with latency to enter an open arm (0.19367 *±* 0.078624, p=0.030). (D-F) There were no differences in percent time spent in open arms between alcohol- and water-exposed controls, but within alcohol-exposed subjects, alcohol consumed during the last binge was negatively associated with time in open arms (−6.9308 *±* 3.1176, p=0.045). (G-I) Similarly, percent open arm entries were similar between alcohol- and water-exposed groups, but last binge alcohol was negatively associated with percent open arm entries (−7.5471 *±* 2.6988, p=0.015). (J-L) No influence of alcohol exposure was detected on the number of open arm entries, but females entered open arms more frequently than males (male: -114.73 *±* 46.288, p=0.028). (M-O) Number of head dips was not dependent upon alcohol exposure variables or sex. (P-R) Subjects that had access to alcohol traveled a shorter distance than water-exposed control subjects (EtOH: -174.94 *±* 77.49, p=0.031), although this did not vary by total or last binge alcohol consumption among alcohol-exposed subjects. For scatterplots, lines of best fit were added independent of the described analysis to aid visualization. * = p<0.05, ** = p<0.01, *** = p<0.001

**Supplementary Figure 3.**
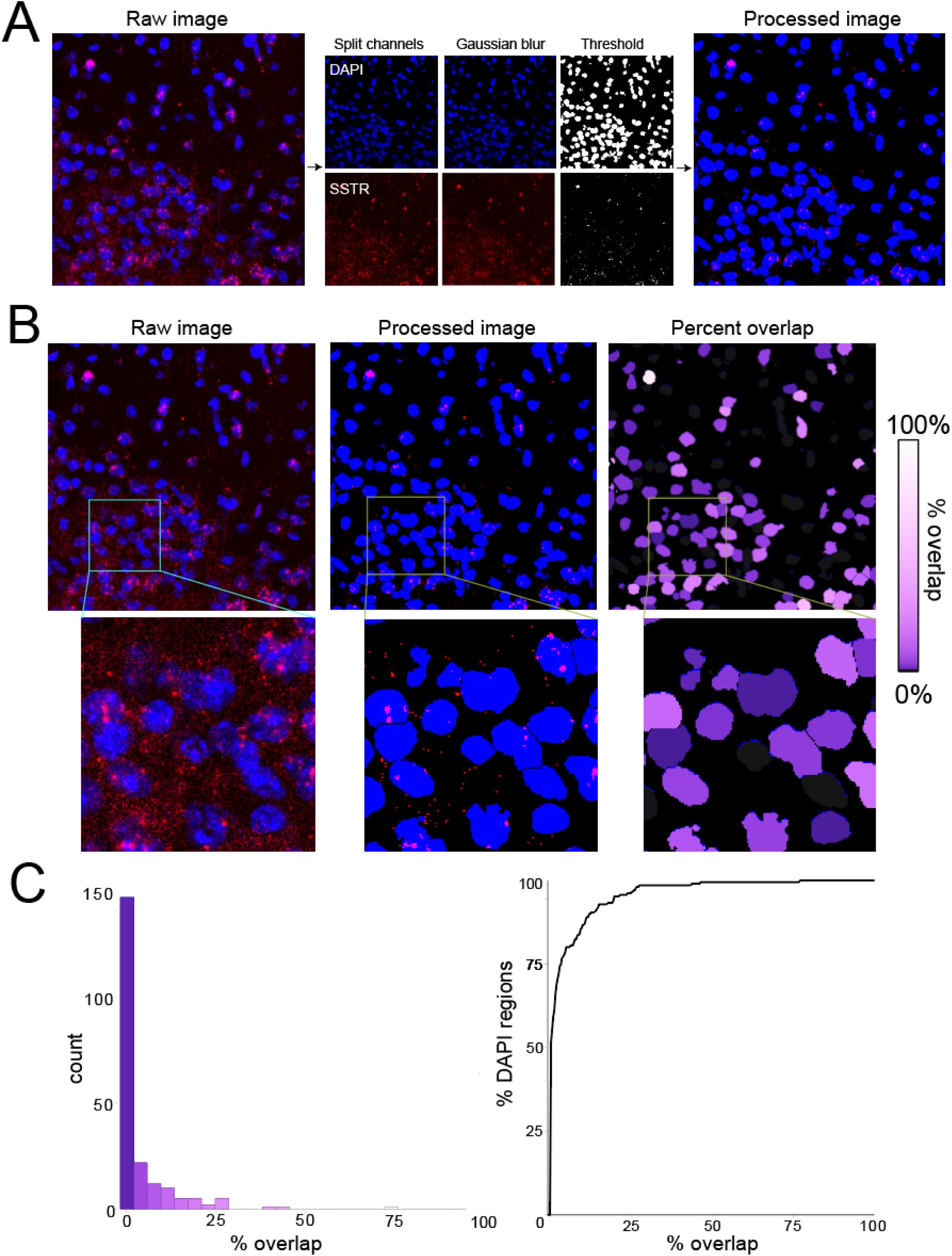
Image processing pipeline. (A) Schematic of the image processing algorithm used to identify DAPI regions and SSTR puncta. (B) Close-ups of raw, processed, and overlap-measured images, with accompanying insets. (C) Histogram and cumulative distribution function of overlap-measured image in B.

**Supplementary Figure 4.**
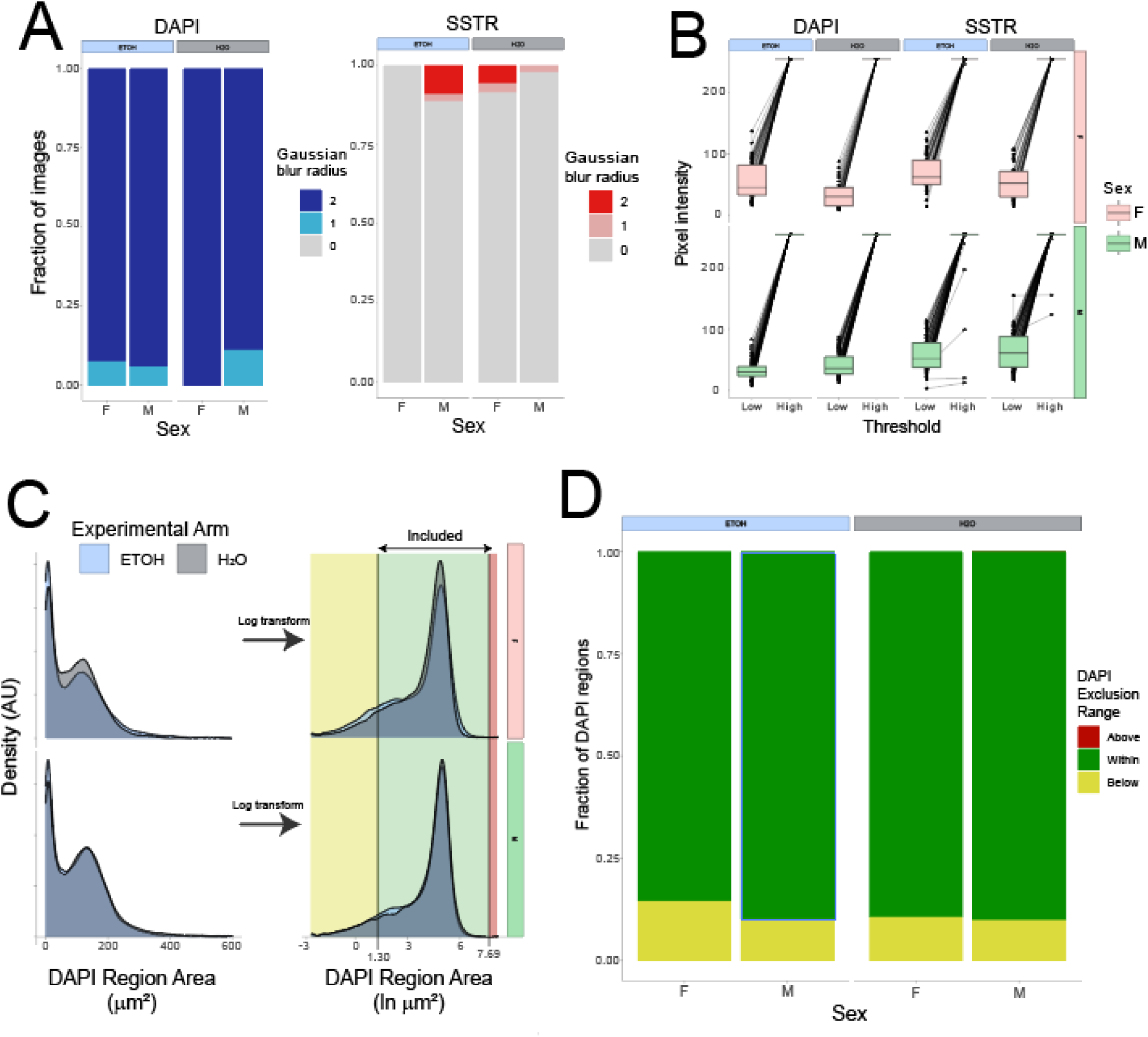
Descriptive statistics from experiment-wise image registration. (A) Manual selection of gaussian blur radius, in both DAPI and SSTR channel0, shown across experimental factors. (B) Manual selection of low and high pixel threshold values for DAPI and SSTR channels, shown across experimental factors. (C) Experiment-wise distribution of DAPI region areas, before and after log-transform, with inclusion range used across the experiment. (D). DAPI region exclusion shown across experimental factors.

## REFERENCES

1. Koob, G.F. and N.D. Volkow, Neurocircuitry of Addiction. Neuropsychopharmacology, 2010/01. 35(1).

2. Kupferschmidt, D.A., et al., *Prefrontal Interneurons: Populations, Pathways,* and Plasticity Supporting Typical and Disordered Cognition in Rodent Models. Journal of Neuroscience, 2022-11-09. 42(45).

3. Brockway, D.F. and N.A. Crowley, *Frontiers | Turning the* ′*Tides on Neuropsychiatric Diseases: The Role of Peptides in the Prefrontal Cortex*. Frontiers in Behavioral Neuroscience, 2020/10/20. 14.

4. Brockway, D., et al., Somatostatin peptide signaling dampens cortical circuits and promotes exploratory behavior. Cell Reports, 2023/08/29. 42(8).

5. Obie Allen, I., et al., Differential Serum Levels of CACNA1C, Circadian Rhythm and Stress Response Molecules in Subjects with Bipolar Disorder: Associations with Genetic and Clinical Factors. medRxiv, 2024.

6. Dao, N.C., et al., Somatostatin neurons control an alcohol binge drinking prelimbic microcircuit in mice. Neuropsychopharmacology 2021 46:11, 2021-06-10. 46(11).

7. Joffe, M.E., D.G. Winder, and P.J. Conn, Contrasting sex-dependent adaptations to synaptic physiology and membrane properties of prefrontal cortex interneuron subtypes in a mouse model of binge drinking. Neuropharmacology, 2020/11/01. 178.

8. NL, H., et al., Effects of acute alcohol on excitability in the CNS - PubMed. Neuropharmacology, 08/01/2017. 122.

9. Lovinger, D.M. and M. Roberto, Synaptic Effects Induced by Alcohol. Current Topics in Behavioral Neurosciences, 2023.

10. Abrahao, K.P., A.G. Salinas, and D.M. Lovinger, Alcohol and the Brain: Neuronal Molecular Targets, Synapses, and Circuits. Neuron, 2017/12/20. 96(6).

11. An integrative theory of prefrontal cortex function - PubMed. Annual review of neuroscience, 2001. 24(1).

12. Carlén, M., What constitutes the prefrontal cortex? Science, 2017. 358(6362): p. 478–482.

13. Volkow, N.D., et al., Addiction: Beyond dopamine reward circuitry. Proceedings of the National Academy of Sciences, 2011-9-13. 108(37).

14. Varodayan, F.P., et al., Morphological and functional evidence of increased excitatory signaling in the prelimbic cortex during ethanol withdrawal. Neuropharmacology, 2018/05/01. 133.

15. Pleil, K.E., et al., Effects of chronic ethanol exposure on neuronal function in the prefrontal cortex and extended amygdala. Neuropharmacology, 2015/12/01. 99.

16. Dao, N.C., et al., Frontiers | Forced Abstinence From Alcohol Induces Sex-Specific Depression-Like Behavioral and Neural Adaptations in Somatostatin Neurons in Cortical and Amygdalar Regions. Frontiers in Behavioral Neuroscience, 2020/05/27. 14.

17. Sicher, A.R., et al., Adolescent binge drinking leads to long-lasting changes in cortical microcircuits in mice. Neuropharmacology, 2023/08/15. 234.

18. Cummings, K.A., et al., Prefrontal somatostatin interneurons encode fear memory. Nature Neuroscience 2019 23:1, 2019-12-16. 23(1).

19. Crowley, N.A. and M.E. Joffe, Developing breakthrough psychiatric treatments by modulating G protein-coupled receptors on prefrontal cortex somatostatin interneurons. Neuropsychopharmacology 2021 47:1, 2021-08-02. 47(1).

20. J, U.-C. and B. AL, Somatostatin-expressing neurons in cortical networks - PubMed. Nature reviews. Neuroscience, 2016 Jul. 17(7).

21. R, O., et al., Patterns of functional connectivity alterations induced by alcohol reflect somatostatin interneuron expression in the human cerebral cortex - PubMed. Scientific reports, 05/12/2022. 12(1).

22. Dao, N., D. Brockway, and C. NA, In Vitro Optogenetic Characterization of Neuropeptide Release from Prefrontal Cortical Somatostatin Neurons. Neuroscience, 2019/11/01. 419.

23. Tripp, A., et al., Reduced somatostatin in subgenual anterior cingulate cortex in major depression. Neurobiology of Disease, 2011/04/01. 42(1).

24. Crowley, N.A. and M.E. Joffe, Developing breakthrough psychiatric treatments by modulating G protein-coupled receptors on prefrontal cortex somatostatin interneurons. Neuropsychopharmacology, 2021 Aug 2. 47(1).

25. JS, R., et al., Evaluation of a simple model of ethanol drinking to intoxication in C57BL/6J mice - PubMed. Physiology & behavior, 01/31/2005. 84(1).

26. Thiele, T.E. and M. Navarro, “Drinking in the Dark” (DID) Procedures: A Model of Binge-Like Ethanol Drinking in Non-Dependent Mice. Alcohol (Fayetteville, N.Y.), 2013 Oct 29. 48(3).

27. Rinker, J.A., et al., Extended Amygdala to Ventral Tegmental Area Corticotropin-Releasing Factor Circuit Controls Binge Ethanol Intake. Biological Psychiatry, 2017/06/01. 81(11).

28. Mathis, A., et al., DeepLabCut: markerless pose estimation of user-defined body parts with deep learning. Nature Neuroscience 2018 21:9, 2018-08-20. 21(9).

29. Al-Hasani, R., et al., Distinct Subpopulations of Nucleus Accumbens Dynorphin Neurons Drive Aversion and Reward. Neuron, 2015/09/02. 87(5).

30. JT, T., et al., Preparation of Acute Brain Slices Using an Optimized N-Methyl-D-glucamine Protective Recovery Method - PubMed. Journal of visualized experiments : JoVE, 02/26/2018(132).

31. Wang, F., et al., RNAscope. The Journal of Molecular Diagnostics, 2012. 14(1): p. 22–29.

32. Lin, L. and E. Sibille, Somatostatin, neuronal vulnerability and behavioral emotionality. Molecular psychiatry, 2015 Jan 20. 20(3).

33. Tomoda, T., et al., Molecular origin of somatostatin-positive neuron vulnerability. Molecular psychiatry, 2022 Feb 10. 27(4).

34. Girgenti, M.J., et al., Prefrontal cortex interneurons display dynamic sex-specific stress-induced transcriptomes. Translational Psychiatry, 2019 Nov 11. 9(1).

35. West, R.K., et al., Recurrent binge ethanol is associated with significant loss of dentate gyrus granule neurons in female rats despite concomitant increase in neurogenesis. Neuropharmacology, 2019/04/01. 148.

36. Catapano, L.A. and H.K. Manji, G Protein-Coupled Receptors in Major Psychiatric Disorders. Biochimica et biophysica acta, 2006 Oct 3. 1768(4).

37. YC, P., Somatostatin and its receptor family - PubMed. Frontiers in neuroendocrinology, 1999 Jul. 20(3).

38. Scheggia, D., et al., Somatostatin interneurons in the prefrontal cortex control affective state discrimination in mice. Nature Neuroscience 2019 23:1, 2019-12-16. 23(1).

39. Park, E., M.B. Mosso, and A.L. Barth, Neocortical somatostatin neuron diversity in cognition and learning. Trends in Neurosciences, 2025. 0(0).

40. Yao, Z., et al., A taxonomy of transcriptomic cell types across the isocortex and hippocampal formation. Cell, 2021/06/10. 184(12).

41. Yao, Z., et al., A high-resolution transcriptomic and spatial atlas of cell types in the whole mouse brain. Nature 2023 624:7991, 2023-12-13. 624(7991).

42. Hostetler, R.E., H. Hu, and A. Agmon, Genetically Defined Subtypes of Somatostatin-Containing Cortical Interneurons. eNeuro, 2023-08-01. 10(8).

43. Cummings, K.A., et al., Control of fear by discrete prefrontal GABAergic populations encoding valence-specific information. Neuron, 2022/09/21. 110(18).

44. Sohn, J., et al., *Preprodynorphin*-*expressing neurons constitute a large subgroup of somatostatin*-*expressing GABAergic interneurons in the mouse neocortex*. Journal of Comparative Neurology, 2014/05/01. 522(7).

45. Wang, H., et al., A tool kit of highly selective and sensitive genetically encoded neuropeptide sensors. Science, 2023-11-17. 382(6672).

